# Chemoselective Halogenation of Premarineosin A for Next-Generation Antimalarial Development

**DOI:** 10.64898/2026.06.16.732709

**Authors:** Natalia R. Harris, Sahar Amin, Brian J. Curtis, Awet A. Teklemichael, Patricia Dranchak, Christina M. McBride, Linnea Verhey-Henke, Chloe J. Warrell, William M. Dulaney, Erin N. Oliphant, James Inglese, Xin-zhuan Su, David H. Sherman, Filipa Pereira

## Abstract

Premarineosin A undergoes rapid, chemoselective C12 halogenation under mild conditions, providing brominated, chlorinated, fluorinated, and iodinated analogs. These derivatives retained potent antiplasmodial activity against both chloroquine-sensitive and -resistant *Plasmodium falciparum* strains and displayed smaller reductions in potency against the resistant strain than the parent compound.

## Introduction

Prodiginines are structurally diverse tripyrrolic natural products that exhibit broad anticancer, antimicrobial, and antiparasitic activities.^1–10^ Among these, premarineosin A (**1**, Figure 1) is a cyclic prodiginine that has potent single-digit nanomolar activity against the malaria parasite *Plasmodium falciparum*.^8,11^ The unique caged macrocyclic architecture of **1** makes it an attractive scaffold for late-stage diversification, particularly through selective functionalization of its tripyrrolic core.

**Figure 1.**
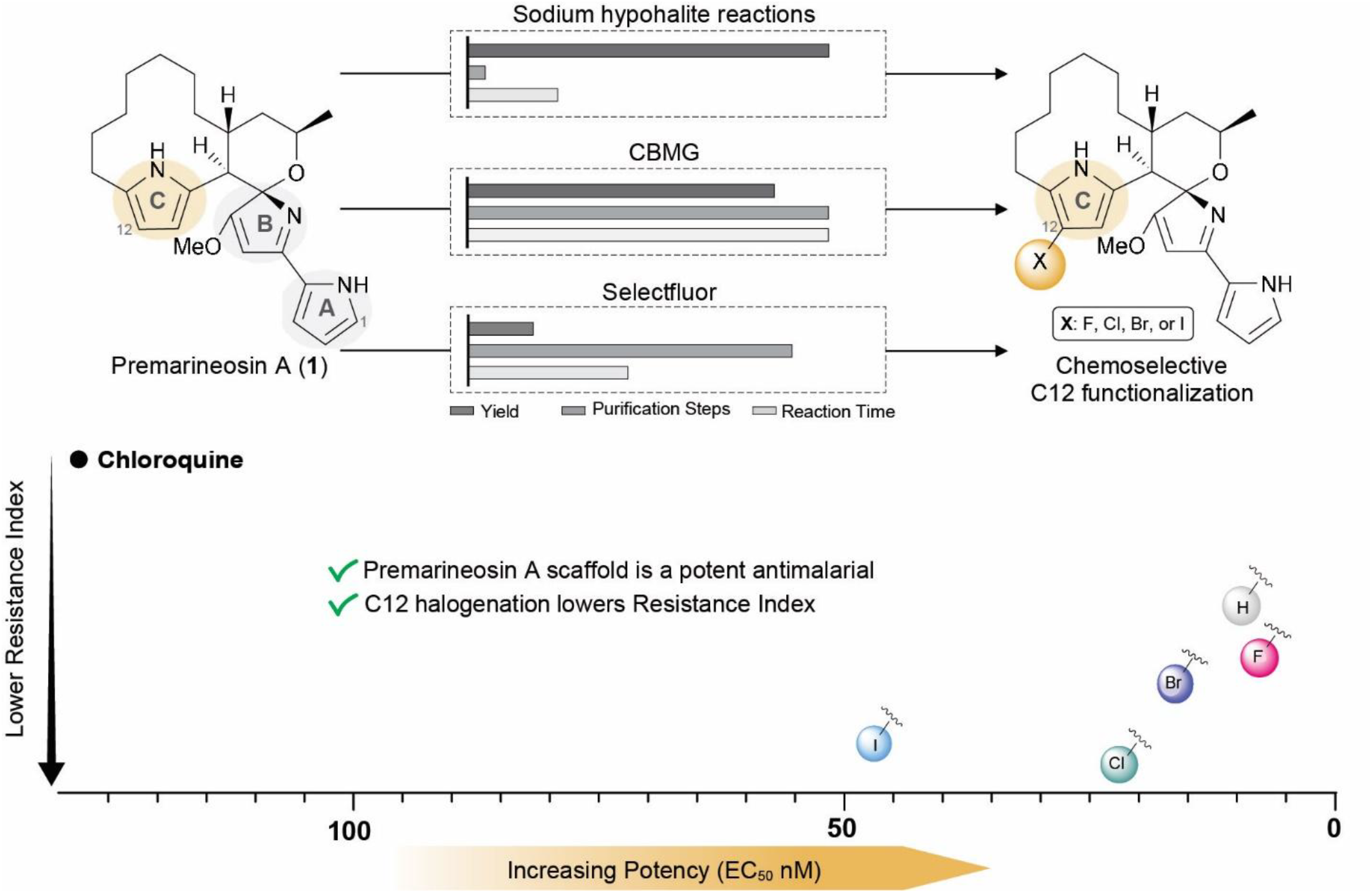
Overview of late-stage C12 halogenation strategies for premarineosin. **A.** Comparative analysis of Selectfluor-, CBMG-, and sodium hypohalite-mediated halogenation reactions highlighting differences in reaction efficiency, purification requirements, and reaction time. Sodium hypohalite conditions enabled rapid and operationally straightforward halogenation, with near-instantaneous conversion following brief mixing and requiring only solvent removal and a single preparative HPLC purification step. All conditions resulted in highly chemoselective functionalization at the C12 position of the C-ring, affording fluoro, chloro, bromo, and iodo premarineosin A derivatives. Halogenated analogs retained potent antimalarial activity and exhibited a lower resistance index relative to chloroquine.

Antimalarial drug resistance remains one of the greatest threats to global malaria control efforts.^12^ Historically, the widespread emergence of resistance to chloroquine and other frontline therapies has repeatedly undermined treatment efficacy and reversed decades of progress in decreasing malaria-associated mortality.^13–15^ Resistance arises through selective pressure, where parasites with reduced susceptibility survive treatment and expand within the parasite population, ultimately leading traditional therapies to fail.^16–18^ Consequently, compounds capable of maintaining potent activity against drug-resistant parasite strains are of particular interest, as they may provide therapeutic advantages in regions where resistance to frontline antimalarial agents has emerged.^19^ Malaria continues to place nearly half of the global population at risk of infection and caused an estimated 282 million cases and over 600,000 deaths in 2024 alone.^12^ As *Plasmodium* parasites continue to evolve resistance to current antimalarial therapies, the development of new agents capable of retaining potency against drug-resistant strains remains increasingly urgent.^12,20^

Halogenation is a valuable late-stage functionalization strategy due to its ability to alter physiochemical and biological properties of small molecules, including modulation of lipophilicity, metabolic stability, and target interactions.^21,22^ Previously, we demonstrated that late-stage bromination of **1** could be achieved through both chemical and enzymatic methods, yielding regioisomeric products with differing biological activities.^11^ While enzymatic bromination of ring A reduced antiparasitic activity, chemical bromination selectively functionalized the C-ring and retained potent activity, identifying this region as a promising site for scaffold diversification.^11^ In particular, carbon 12 (C12) emerged as a site of interest for further derivatization and expansion to additional halogen substituents (Figure 1). Initial access to 12-bromo premarineosin A was achieved via N-bromosuccinimide (NBS); however, the reaction resulted in a moderate yield (55%), highlighting the need for more efficient strategies to access a broader range of C12-halogenated analogs.^11^

In this work, we expanded our investigation of late-stage electrophilic halogenation strategies of **1** under operationally facile, mild conditions that proceed rapidly and with improved efficiency. These transformations provided streamlined access to a series of C12-halogenated analogs that retained potent activity against both chloroquine-sensitive and chloroquine-resistant *P. falciparum* strains (Figure 1). Collectively, these findings demonstrate that strategic late-stage halogenation of premarineosin A provides an effective route to next-generation antimalarial analogs capable of maintaining strong antiparasitic activity in a clinically relevant drug-resistant parasite background.

## Results and Discussion

### Halogenation of premarineosin A at C12 position

Fluorine substitution is among the most widely used strategies in medicinal chemistry for modulating bioactivity while minimizing perturbations to molecular size or steric interactions.^23,24^ Due to its small atomic radius and high electronegativity, fluorine often serves as a bioisosteric replacement, preserving the scaffold while altering electronic properties to improve drug-like behavior.^23–25^ Treatment of **1** with Selectfluor^26^ generated 12-fluoro premarineosin A (**2**) in moderate yield, confirmed via NMR and LC-MS/MS (20%, Figure 2A, Supplementary Figures 1A & C, 2A, 3, & 9-15).

**Figure 2.**
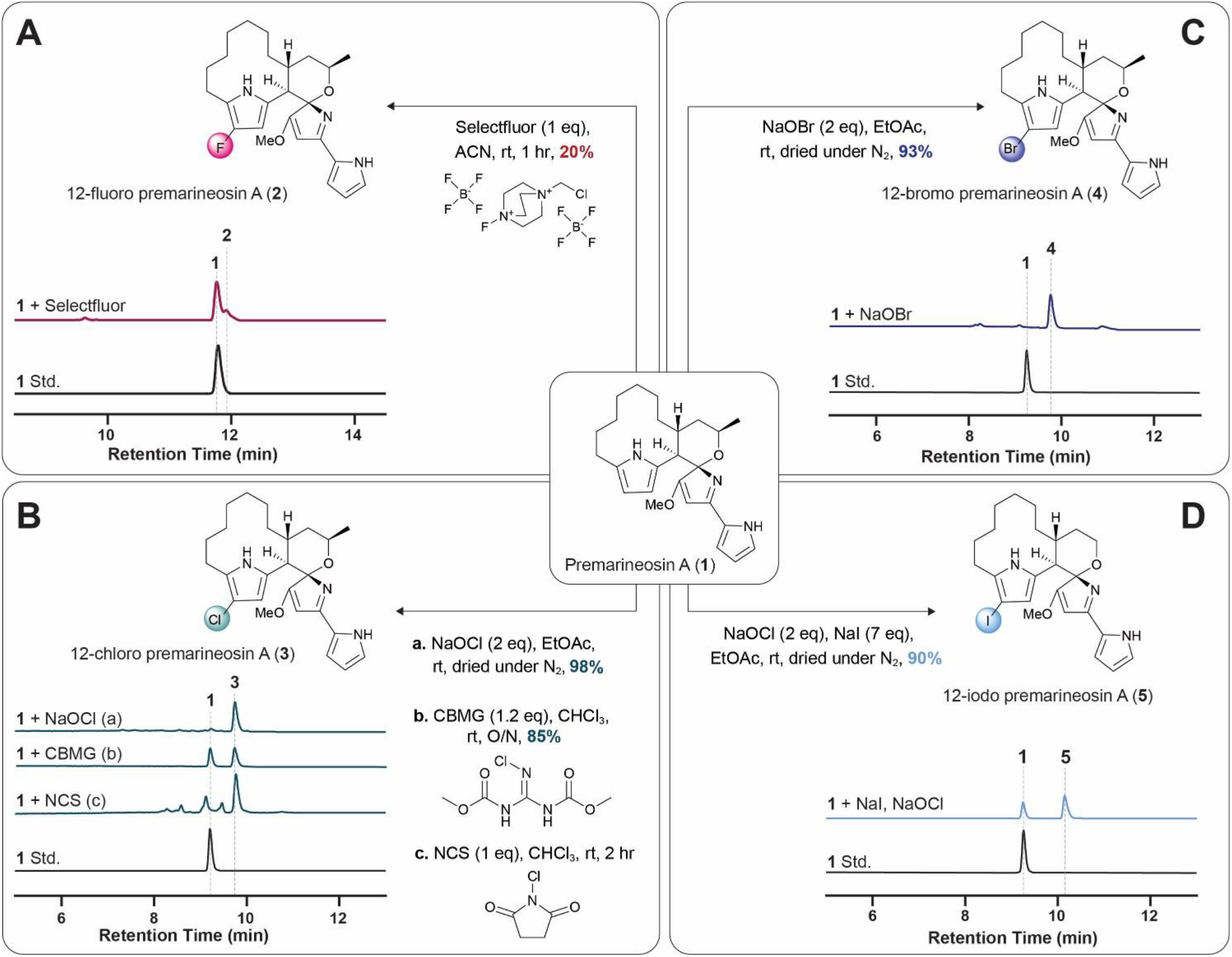
Development of late-stage halogenation reactions for premarineosin. **A.** Reaction optimization and HPLC analysis of fluorination, chlorination, bromination, and iodination conditions targeting selective C12 functionalization of the premarineosin A (**1**) scaffold. (**A**) Treatment with Selectfluor afforded 12-fluoro premarineosin A (**2**) in moderate yield following a 1 hr reaction in acetonitrile. (**B**) Chlorination studies demonstrated substantial differences in efficiency depending on the halogenating reagent employed. Sodium hypochlorite-mediated chlorination (a) rapidly generated 12-chloro premarineosin A (**3**) under mild conditions at 98% yield, whereas CBMG required overnight incubation to achieve partial conversion (b) and NCS resulted in substantial decomposition and formation of multiple by-products (c). (**C**) Bromination using sodium hypobromite in ethyl acetate proceeded cleanly and rapidly to afford 12-bromo premarineosin A (**4**) in 93% yield with minimal detectable degradation. (**D**) Treatment with sodium iodide and sodium hypochlorite generated 12-iodo premarineosin A (**5**) in 90% yield. HPLC chromatograms illustrate selective conversion of **1** to the corresponding mono-halogenated products under each reaction condition.

We next explored complementary late-stage chlorination using N-chlorosuccinimide (NCS)^27^, 2-chloro-1,3-bis(methoxycarbonyl)guanidine (CBMG)^28^, and sodium hypochlorite (NaOCl). Initial efforts using NCS resulted in moderate conversion with substantial degradation of **1**, mirroring the challenges previously encountered with NBS-mediated bromination reactions (Figure 2B, Supplemental Figure 1A & D).^11^ In contrast, chlorination with CBMG produced 12-chloro-premarinosin A (**3**) with improved selectivity, although the reaction required overnight incubation and reached only partial conversion to product (Figure 2B, Supplementary Figures 1A & D, 16-20). Treatment with sodium hypochlorite (2 equiv.) proved significantly more efficient, resulting in immediate and near-quantitative conversion to the chlorinated analog (98%, Figure 2B, Supplementary Figures 1A & D, 2B, 3, & 21-23).^29,30^ Unlike the Selectfluor and CBMG reactions, which required extended reaction times, aqueous quenching, extraction, and multistep purification, the sodium hypochlorite-mediated reaction proceeded almost instantaneously and required only brief mixing, solvent removal, and a single preparative HPLC purification step. Importantly, all chlorination strategies maintained regioselectivity for C12 functionalization despite differences in efficiency and reaction rate.

Encouraged by the efficiency of the sodium hypochlorite-mediated chlorination, we next applied analogous sodium hypohalite conditions to bromination and iodination reactions. Bromination proved particularly efficient. Treatment of **1** with sodium hypobromite in ethyl acetate resulted in immediate and near-complete conversion to a single brominated product (**4**), which was isolated in 93% yield (Figure 2C, Supplementary Figures 1A & E, 2C, 3).^29,30^ Similar to the sodium hypochlorite-mediated chlorination, the reaction was operationally straightforward, requiring only brief mixing prior to solvent removal and direct purification by preparative HPLC. In contrast to our previous NBS-mediated strategy, the reactions proceeded cleanly with minimal detectable decomposition of the starting material. Structural analysis via NMR and LC-MS/MS confirmed regioselective bromination at C12 (Supplementary Figures 24-26).^11^

Using similar mild conditions, iodination with sodium iodide (NaI) and sodium hypochlorite rapidly generated 12-iodo premarineosin A (**5**) in 90% yield (Figure 2D, Supplementary Figures 1A & E, 2D, 3, & 27-33).^30^ The NaOCl/NaI system likewise proceeded instantaneously and avoided the extended reaction times, quenching procedures, and extraction workflows required for Selectfluor- and CBMG-mediated halogenation reactions.

Halogenation of pyrrolic natural products is often complicated by nonselective halogenation and scaffold decomposition, frequently requiring protecting group strategies or extensive optimization.^31–33^ In contrast, these reactions proceed rapidly under mild conditions with inexpensive and operationally convenient reagents, enabling high-yielding late-stage diversification of the premarineosin A (**1**) scaffold. Notably, despite the well-established preference for electrophilic functionalization at unsubstituted α-positions of pyrroles, all reactions proceeded selectively at the C12 (β) position of the C-ring.^34^ While unsubstituted α-carbons are generally the preferred sites for pyrrolic C–H functionalization, suggesting that halogenation at the C1-position (A-ring, Figure 1) might be favored, all tested conditions instead resulted in highly selective halogenation at the C12 position (C-ring, Figure 1). We reason that the unique conformational and electronic properties of the caged macrocyclic architecture preferentially activate the C-ring toward electrophilic aromatic substitution under these conditions.

### Bioactivity assays

Evaluation of the halogenated premarineosin A analogs against both chloroquine-sensitive (3D7) and chloroquine-resistant (Dd2) *P. falciparum* strains revealed that C12 halogenation was generally associated with smaller potency differences between the two parasite strains relative to the parent scaffold (Figure 3, Supplementary Figure 4). Premarineosin A (**1**) displayed potent activity against the 3D7 strain (EC_50_ = 2.0 ± 1.5 nM), but potency was reduced against the resistant Dd2 strain, resulting in a resistance index (RI) of 4.4. All halogenated analogs exhibited a lower resistance index than premarineosin A, suggesting that their antiplasmodial activity was less affected by the chloroquine-resistant Dd2 genetic background than the parent compound. Among the series, 12-chloro premarineosin A (**3**) showed the greatest improvement, with nearly equivalent activity across both strains (RI = 1.2), despite a modest reduction in absolute potency (Figure 3). Similarly, the iodinated and brominated analogs exhibited improved resistance indices of 1.4 and 2.3, respectively. Notably, 12-fluoro premarineosin A (**2**) showed a reduced RI (2.9) while maintaining the low-nanomolar potency observed for the parental compound. All analogs maintained favorable selectivity indices (SI) ranging from 30.8 to 445.0 (Figure 3, Supplementary Figure 5). Overall, these results demonstrate that late-stage C12 halogenation provides an effective strategy to modulate the biological activity of the premarineosin scaffold while retaining potent antiplasmodial activity against both chloroquine-sensitive and chloroquine-resistant parasite strains.

**Figure 3.**
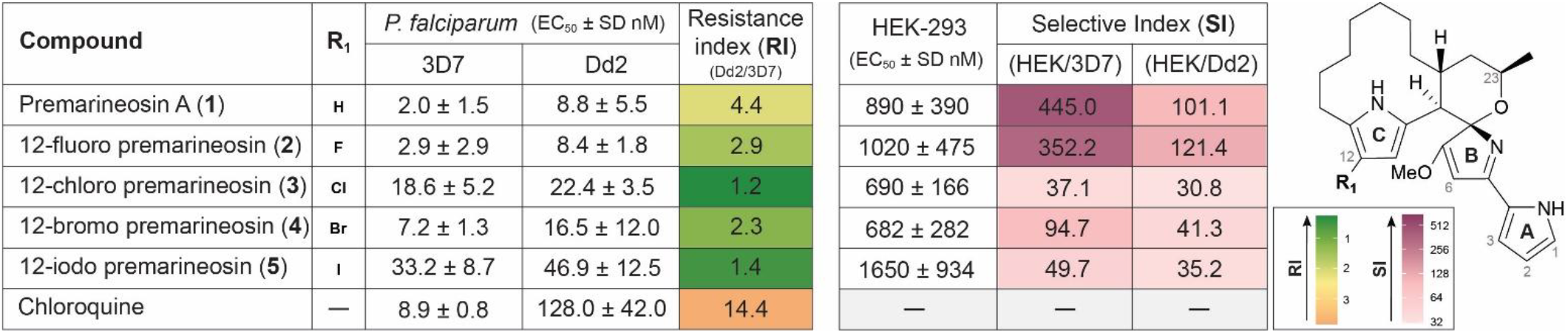
Antiplasmodial activity, resistance indices, and mammalian cytotoxicity profiling of halogenated premarineosin A derivatives. In vitro antiplasmodial activity of premarineosin A and C12-halogenated analogs against drug-sensitive *Plasmodium falciparum* 3D7 and chloroquine-resistant Dd2 parasite strains. Resistance index (RI) values were calculated as the ratio of Dd2 to 3D7 EC_50_ values. C12 halogenation resulted in substantially lowered resistance index relative to the parent scaffold and chloroquine, with the chloro- and iodo-substituted analogs displaying the lowest RI values. Cytotoxicity was evaluated in HEK-293 cells, and selective indices (SI) were calculated relative to antiplasmodial activity against 3D7 and Dd2 parasites. While fluorinated and brominated analogs retained favorable selectivity profiles, chlorinated, and iodinated derivatives displayed reduced selectivity despite improved resistance profiles. Heat map coloring represents relative RI and SI values across the compound series. Values are reported as mean EC_50_ ± SD from six and three biological replicates for cytotoxicity and antimalarial assays, respectively.

## Conclusion

In summary, we demonstrated that premarineosin A undergoes rapid and selective C12 halogenation under mild conditions, enabling efficient access to a diverse series of semisynthetic analogs with improved chemoselectivity and synthetic efficiency. Importantly, C12 halogenation preserved potent antiparasitic activity across both chloroquine-sensitive and chloroquine-resistant *P. falciparum* strains, with several analogs exhibiting smaller potency shifts in the Dd2 background than the parent compound. As *Plasmodium* parasites continue to evolve resistance to current antimalarial agents, these findings establish strategic late-stage halogenation of premarineosin A as a promising approach for generating antimalarial analogs that retain potent activity against both drug-sensitive and drug-resistant parasite strains.

## Materials and Methods

Reagents and solvents were purchased from EMD Millipore, Sigma-Aldrich, Oakwood Chemical, Chem Impex, Thermo-Fisher Scientific, AABlocks, Advanced Chem Blocks, TCI, or Arctom unless indicated otherwise. NMR spectra were recorded on a Bruker 600 NMR system (600 MHz). High-resolution mass spectra and analytical reaction analysis were recorded on an Agilent Technologies 6250 TOF LC/MS equipped with an Agilent 1290 Infinity II HPLC and analytical HPLC (Shimadzu) equipped with a PDA detector. Samples were also analyzed via ultra-high-performance liquid chromatography-quadrupole time-of-flight mass spectrometry (UHPLC-LCMS) on an Agilent 1290 Infinity II UHPLC coupled to an Agilent 6545 ESI-Q-TOF-MS.

### Premarineosin A production, isolation, and purification

Premarineosin A was produced, isolated, and purified as previously described.^11^ Briefly, strains of *Streptomyces eitanensis* engineered for overproduction of premarineosin A were cultured in 1L producing media (GICYE: 10 g/L glucose, 30 g/L inulin, 5 g/L bacto peptone, 10 g/L corn gluten meal, 5 g/L yeast extract, 2 g/L CaCO_3_ in ultrapure water) in 2.8 L Fernbach flasks for seven days at 22 °C and 165 rpm. After seven days, the total cell biomass was collected by vacuum filtration and extracted overnight in 1 L of acetone with shaking. The acetone was retained as filtrate, concentrated, and liquid-liquid extracted with ethyl acetate. The resulting crude organic extract was dried-loaded onto silica gel and purified by normal-phase flash chromatography on a Biotage Isolera system using prepacked 40 g silica gel cartridges (SiliaSep). Initial elutions utilized an ethyl acetate/hexane gradient (15-100%) with 1% acetic acid; premarineosin A eluted at 34-50%. Premarineosin A-containing fractions were further purified using a 3% ammonia (7N in methanol) in DCM solution / hexane gradient (10-50%); premarineosin A eluted at 20-30%. To obtain high purity (>95%) premarineosin A with a protonation state matching that of the halogenated derivatives, premarineosin A fractions were purified using preparative HPLC with a gradient of acetonitrile and water, both modified with 0.1% formic acid. Premarineosin purified under basic conditions retained potent antiplasmodial activity against *Plasmodium falciparum* Dd2, with an IC_50_ of 2.6 ± 1.1 nM, consistent with our previous reports.^11^ In contrast, premarineosin purified under acidic conditions (consistent with halogenated analog production) displayed reduced, though still potent, activity with an IC_50_ of 8.8 ± 5.5 nM against *P. falciparum* Dd2. Structure and purity were confirmed via LC-MS, LC-MS/MS and 2-D NMR.

### Bromination of premarineosin

Premarineosin A (20 µL, 100 mM stock solution, 1.0 equiv) was added to ethyl acetate (5 mL). Sodium hypobromite (2.0 equiv) was then introduced to initiate bromination.^29,30^ The reaction mixture was briefly mixed, and the solvent was immediately dried under nitrogen. LC-MS (Supplementary Figure 1) and HPLC (Figure 2C) analysis of the crude residue revealed near-quantitative conversion (∼100%) to a single brominated product. For large scale, the reactions were conducted ten-fold and purified via preparative HPLC per the methods below. Structural characterization was confirmed via LC-MS, LC-MS/MS and 2-D NMR.

### Iodination of premarineosin A

Premarineosin A (20 µL, 100 mM stock solution, 1.0 equiv) was added to ethyl acetate (5 mL) followed by sodium iodide (20 µL, 2 M aqueous solution, 20 equiv).^29,30^ The reaction was initiated by addition of sodium hypochlorite (20 µL, ∼7 equiv) and stirred at room temperature for 5 min. Reaction was stopped by drying down under nitrogen. LC-MS (Supplementary Figure 1) and HPLC (Figure 2D) analysis of the crude residue revealed ∼90% conversion to a single iodinated product. For large scale, the reactions were conducted ten-fold and purified via preparative HPLC per the methods below. Structural characterization was confirmed via LC-MS, LC-MS/MS and 2-D NMR.

### Chlorination of premarineosin A

Reaction with CBMG (small scale): A solution of premarineosin A (10 μL, 10 mM stock solution) was diluted with solvent (5 μL) in a reaction vial. A solution of 2-Chloro-1,3-bis(methoxycarbonyl)guanidine (CBMG) (5 μL, 1.2 equiv, 30 mM stock solution) in chloroform was then added to give a reaction volume of 20 μL and allowed to stir overnight at room temperature.^28^ Reaction was analyzed using LC-MS (Supplementary Figure 1) and HPLC (Figure 2B).

Reaction with CBMG (large scale): To a solution of premarineosin A (10 mg, 0.025 mmol) in chloroform under stirring, a solution of 2-Chloro-1,3-bis(methoxycarbonyl)guanidine (CBMG) (7.7 mg, 0.029 mmol, 1.2 equiv) in chloroform was added dropwise and left overnight.^28^ The reaction mixture was monitored by thin layer chromatography (7 % MeOH in DCM containing 3% concentrated NH_3_ (v/v)) until completion, where the mixture was concentrated. The crude mixture was purified using reverse-phase flash column chromatography on a C18 cartridge using water/acetonitrile gradient followed by preparative HPLC per the methods below. Structural characterization was confirmed via LC-MS, LC-MS/MS and 2-D NMR. Reaction with NaOCl: Premarineosin A (20 µL, 100 mM stock solution, 1.0 equiv) was added to ethyl acetate (5 mL).^29,30^ Sodium hypochlorite (2.0 equiv) was then introduced to initiate chlorination. The reaction mixture was briefly mixed, and the solvent was immediately dried under air. LC–MS (Supplementary Figure 1) analysis of the crude residue revealed near-quantitative conversion (∼100%) to a single chlorinated product. For large scale, the reactions were conducted ten-fold and purified via preparative HPLC (Figure 2B) per the methods below. Structural characterization was confirmed via LC-MS, LC-MS/MS and 2-D NMR.

Reaction with NCS: To a solution of premarineosin A (1mg, 0.025 mmol) in chloroform under stirring, a solution of NCS (0.3mg, 0.025 mmol, 1 equiv) in chloroform was added dropwise.^27^ After two hours, the reaction mixture was quenched with saturated sodium bicarbonate solution and the organic layer was extracted. Reaction was analyzed using LC-MS (Supplementary Figure 1) and HPLC (Figure 2B).

### Fluorination of premarineosin A

Reaction with Selectfluor (small scale): A solution of premarineosin A (10 μL, 10 mM stock solution) was diluted with solvent (6.7 μL) in a reaction vial. A stock solution of Selectfluor (3.3 μL, 30 mM stock solution, 1 equiv) in acetonitrile was then added to give a reaction volume of 20 μL.^26^ After stirring for 1 hour at room temperature, saturated aqueous sodium bicarbonate was added to quench the reaction. The mixture was extracted with dichloromethane (DCM), and the organic layer was dried under a stream of nitrogen. Reaction was analyzed using LC-MS (Supplementary Figure 1) and HPLC (Figure 2A).

Large scale: To a solution of premarineosin A (5 mg, 5 mM stock solution, 1 equiv) in acetonitrile (1.25 mL) under stirring, a solution of Selectfluor (4 mg, 30 mM stock solution, 1 equiv) in acetonitrile (1.0 mL) was added dropwise after complete dissolution. The reaction mixture was diluted with an additional 250 µL of acetonitrile to bring the total volume to 2.5 mL. After stirring for 1 hour at room temperature, saturated aqueous sodium bicarbonate (∼1 mL) was added to quench the reaction.^26^ The mixture was extracted with dichloromethane (DCM), and the organic layer was dried under a stream of nitrogen. The reaction was performed five times on a 5 mg scale and the crude reaction mixtures were combined prior to purification. Purification was achieved using reverse-phase flash chromatography on a C18 cartridge using water/acetonitrile gradient followed by preparative HPLC per the methods below. Structural characterization was confirmed via LC-MS, LC-MS/MS and 2-D NMR.

### TOF-LC-MS Analysis

Reactions were subjected to LC-MS analysis using an Agilent G6230B TOF mass spectrometer operating in positive ESI mode (200–1200 *m/z*). The LC separation was performed using a Kinetex 1.7 μm Phenyl-Hexyl 100 Å LC Column (50 × 2.1 mm) at a flow rate of 0.4 mL/min, with a gradient of 5–40% solvent B (95% acetonitrile + 5% water + 0.1% formic acid) in solvent A (water + 0.1% formic acid) for the first 2 minutes, an isocratic hold at 40% for 5 minutes, following at 40-100% gradient for 3 minutes. The first minute of flow was diverted to waste.

### QTOF-LC-MS/MS Analysis

Purified products were analyzed using ultra-high-performance liquid chromatography coupled with quadrupole time-of-flight mass spectrometry (UHPLC-LCMS) performed with an Agilent 1290 Infinity II UHPLC coupled to an Agilent 6545 ESI-Q-TOF-MS. The samples were injected, and data was collected using auto-MS/MS in positive mode. The chromatographic separation was carried out on a Phenomenex Kinetex Phenyl-Hexyl column (1.7 μm, 2.1 × 100 mm). Isocratic elution was performed with 90% solvent A (water + 0.1% formic acid) for 1 min, followed by a 9 min linear gradient to 100% solvent B (95% acetonitrile + 5% water + 0.1% formic acid). Electrospray ionization was performed at a capillary temperature of 320 °C, with a source voltage of 3.5 kV, and a sheath gas flow rate of 11 L/min.

### Analytical HPLC Analysis

Reaction supernatants were analyzed in comparison to standards using an analytical HPLC (Shimadzu) equipped with a PDA detector and analyzed with a Phenyl-Hexyl column (Luna 5 μM Phenyl-Hexyl 100 Å, LC Column 250 × 4.6 mm, RT). Gradient elution was performed with 20% solvent B (acetonitrile + 0.1% formic acid) in solvent A (Water + 0.1% formic acid) to 80% solvent B for the mobile phase at 1 mL/min^-1^ for bromo-, chloro-, and iodination reactions. Gradient elution was performed with 20% solvent B (acetonitrile + 0.1% formic acid) in solvent A (Water + 0.1% formic acid) to 60% solvent B for the mobile phase at 1 mL/min^-1^ for fluorination reactions.

### Reverse-Phase Flash Column Chromatography

The crude reaction mixtures from the fluorination (Selectfluor) and chlorination (CBMG) of premarineosin A were resuspended in 90:10 acetonitrile/water and purified by reversed-phase column chromatography (Biotage) on a C18 cartridge (SiliaSep 12g-C18), using a gradient of 10-80% water/acetonitrile for the fluorinated mixture and 10-40% water/acetonitrile for chlorinated mixture, both performed at a flow rate of 15 mL/min. Products were analyzed using LC-MS, and further purified by preparative HPLC (Shimadzu).

### Preparatory HPLC Purification

The resulting synthetic halogenation material was resuspended in acetonitrile at ∼10-20mg/mL and purified by preparative HPLC (Shimadzu) equipped with a PDA detector using a Phenyl-Hexyl column (5 μm, 100 Å, 250 × 10 mm) and a gradient (specified for each product below) of acetonitrile and water, both modified with 0.1% formic acid, over 100 min at a flow rate of 10 mL min^−1^. Isolated products were analyzed using LC-MS (Supplementary 2) and LC-MS/MS (Supplementary Figures, HPLC (Supplementary Figure 3), and NMR (Supplementary Figures 6-31).

#### 12-fluoro premarineosin A

- 10-37% ACN, 30 min
- 37% ACN hold, 30 min
- 45%-60% ACN, 20 min
- 60%-80 ACN, 20 min
- 80-100% ACN, 20 min

#### 12-chloro premarineosin A

- 10-40% ACN, 30 min
- 40% ACN hold, 30 min
- 45%-60% ACN, 20 min
- 60%-80 ACN, 20 min
- 80-100% ACN, 20 min

#### 12-bromo premarineosin A & 12-iodo premarineosin A

- 10-45% ACN, 30 min
- 45% ACN hold, 30 min
- 45%-80% ACN, 20 min
- 80-100% ACN, 20 min

### CellTiter-Glo (CTG) cell viability assay

A CTG assay (Promega, cat # G7572) was utilized to quantify cellular ATP and evaluate cell viability according to manufacturer protocols and as previously described.^11^ The HEK293 human embryonic kidney cell line was obtained from ATCC (cat # CRL-1573) and was regularly evaluated for Mycoplasma contamination using the MycoAlert PLUS *Mycoplasma* Detection Kit (Lonza Bioscience, cat # LT07) following manufacturer protocols. Cells were maintained in Dulbecco’s Modified Eagle Medium (DMEM) high glucose medium (Gibco, Cat. #11965-092) supplemented with 10% HyClone Characterized Fetal Bovine Serum (Cytiva, cat # SH30071.03) and 100 U/mL penicillin-streptomycin (Gibco, cat # 15140122) and were cultured at 37C with 5% CO_2_ and 95% relative humidity. Cell number and viability were measured on the Countess 3 FL Automated Cell Counter (Invitrogen) using Trypan blue stain (Invitrogen, cat # T10282). A ThermoFisher Multidrop Combi liquid dispenser dispensed 4uL/well cells into white 1536-well microplates (Corning, cat # 7464) at a density of 1500 cells/well and cells were incubated overnight. Evaluated compounds were transferred into respective wells of each plate at 25 nL/well with the Mosquito (SPT Labtech) in 16-pt, 1:3 titrations for a final concentration of 62.5 µM–4.36 pM (initial concentration 10mM). Controls were transferred into respective wells of each plate at 25 nL/well with the mosquito with a 16-pt, 1:2 titration of Digitonin for a final concentration of 125 µM–3.81 nM. The final concentration of DMSO was 0.58%. Cells were incubated for 72 hours before 3 µL/well of CTG was transferred to each plate using the BioRaptr 2.0 FRD (LetsGoRobotics). Plates were incubated in the dark at ambient temperature for 10 min. Luminescence was measured using a ViewLux 1430Ultra HTS (PerkinElmer) with the following optical settings: exposure = 1 s, gain = high, speed = slow, binning = 2X. Dimethyl sulfoxide (“DMSO”; AMRESCO, cat # RGE-3070) was used as a vehicle control. Digitonin (Sigma-Aldrich, cat # D141) was prepared as a 20 mM stock solution in DMSO and stored at −30 °C for use as a cytotoxicity control. Premarineosin A and derivatives were prepared in 10 mM stock solutions in DMSO. Data were normalized to 125 µM Digitonin as −100% inhibition in a row-wise manner across the plate. Normalization was performed in Excel (Microsoft) and Concentration Response Curves (CRCs) were plotted in GraphPad Prism (GraphPad Software, Inc.) with error bars representing the standard deviation of six replicate wells.

### In vitro antimalarial assay.^35^

The antimalarial growth inhibition assay was conducted as previously described.^11^ Briefly, the *Plasmodium falciparum* parasites (strains Dd2 and 3D7) were diluted to 0.75% parasitemia with 2% hematocrit, and 50 μL diluted parasites were added to each well in a 96-well plate containing 50 μL of diluted drug. Compounds were diluted three-fold in triplicate with concentrations ranging from 20 –0.001 µM/10 – 0.0005 µM. The parasites were incubated with the drugs at 37 °C under mixed gas (5% O_2_, 5% CO_2_, and 90% N_2_) conditions for 48 hours. DNA was released from the parasites and stained with SYBR green dye. The plate was set in the dark with gentle agitation for 1 h, and the resulting signals were evaluated in a FLUOstar OPTIMA microplate reader (BMG Labtech, Germany). Each *in vitro* experiment was conducted twice and in triplicate wells. Data were analyzed and plotted using Prism 9.0 software (GraphPad Software, Inc., San Diego, CA).

## Acknowledgements

We are grateful for support from NIH grant R35GM118101 and the Hans W. Vahlteich Professorship (to D.H.S.), a National Science Foundation Graduate Research Fellowship #DGE2241144 (to C.M.M.), the University of Michigan College of Pharmacy Duellman Graduate Student Research Fund #PGG030232 (to S.A.), and a Michigan Pioneer Fellowship (to B.J.C.). This research was supported [in part] by the Intramural Research Program of the National Institutes of Health (NIH). The contributions of the NIH authors are considered Works of the United States Government. The findings and conclusions presented in this paper are those of the author(s) and do not necessarily reflect the views of the NIH or the U.S. Department of Health and Human Services. This research was also supported in part by the Intramural Research Programs of the National Center for Advancing Translational Sciences, NIH under project 1ZIA TR000495-01 (to J.I.). This work was also supported by the University of Michigan BioNMR Core Facility (U-M BioNMR). The U-M BioNMR Core is grateful for support from U-M including the College of Literature, Sciences and Arts, the Life Sciences Institute, the College of Pharmacy, and the U-M Biosciences Initiative. We thank Abigail Davis (NCATS, NIH) for assistance with cell culture and data analysis.

